# Prediction of Neurocognitive Profiles in Youth From Resting State fMRI

**DOI:** 10.1101/495267

**Authors:** Chandra Sripada, Saige Rutherford, Mike Angstadt, Wesley K. Thompson, Monica Luciana, Alex Weigard, Luke Hyde, Mary Heitzeg

## Abstract

Difficulties with higher-order cognitive functions in youth are a potentially important vulnerability factor for the emergence of problematic behaviors and a range of psychopathologies. This study examined 2,013 9-10 year olds in the first data release from the Adolescent Brain Cognitive Development 21-site consortium study in order to identify resting state functional connectivity patterns that predict individual-differences in three domains of higher-order cognitive functions: General Ability, Speed/Flexibility, and Learning/Memory. We found that connectivity patterns involving task control networks and default mode network were prominently implicated in predicting individual differences across participants across all three domains. In addition, for General Ability scores specifically, we observed consistent cross-site generalizability, with statistically significant predictions in 14 out of 15 held-out sites. These findings demonstrate that resting state connectivity can be leveraged to produce generalizable markers of neurocognitive functioning. Additionally, they highlight the importance of task control-default mode network inter-connections as a major locus of individual differences in cognitive functioning in early adolescence.

## Main

Adolescence is a time of major physical, cognitive, and psychosocial change. It is also a time of great vulnerability for the emergence of maladaptive behavioral patterns and psychopathologies (1), which can cascade into poorer mental and physical health throughout adulthood. A major task for clinical neuroscience is to identify pre-morbid features that place the child at elevated risk for future adverse outcomes.

One major risk factor for adolescent psychopathology is difficulty with higher-order cognitive functions, which encompass a diverse set of abilities for reasoning and problem solving, cognitive control and mental flexibility, and learning and recalling information (2–5). Difficulties with higher-order cognitive functions have been associated with a number of disorders, including both externalizing-spectrum disorders (e.g., substance use disorders and attention-deficit/hyperactivity disorder) (6–8) as well internalizing disorders (e.g., depression and anxiety) (9–11). The generality of these associations has led to the interesting suggestion that deficits in some kinds of higher-order cognitive functions (e.g., cognitive control) represent domain-general vulnerability factors for psychopathology (12–14).

Concurrently, there is great interest in the neural underpinnings of higher-order cognitive functions (15). Emerging models focus on interrelationships between networks involved in active control of task processing (16, 17) and default mode network (DMN), involved in spontaneous cognition (18). Task control networks include frontoparietal network (FPN) and cingulo-opercular (CO) network, both involved in top-down control (19, 20), as well as dorsal attention network (DAN), involved in goal-directed attention (21). DMN—active in task-free states and implicated in spontaneous thought, evaluation, and memory (18, 22)—works in both cooperative and antagonistic ways with task control networks (23–26). The salience network (SAL) and ventral attention network (VAN), important for detection of unexpected events and ongoing monitoring (27), play key roles in identifying when adaptive control is required (28, 29). Higher-order cognitive functions are thought to rely on complex interplay between this set of networks, in which task control networks (which we define to include FPN, CO, DAN, SAL, and VAN; see 38) supply task-relevant top-down regulatory signals that modulate spontaneous processing unfolding in association cortices within DMN (16, 31).

Importantly, from early childhood to young adulthood, there is extensive development of functional interconnections between task control networks and DMN (32–34), suggestive of growing informational exchange and maturing top-down regulatory relationships. These observations raise an intriguing question about whether altered connectivity patterns among these networks during youth are predictive of differences in higher-order cognitive functioning.

The present study investigates this question leveraging the first data release from the Adolescent Brain and Cognitive Development (ABCD) national consortium study, which will comprehensively characterize a cohort of over 11,000 adolescents using behavioral, psychosocial, and neuroimaging measures over the course of 10 years. To accomplish this goal, data are collected from 21 sites nationwide. The baseline assessment of the study cohort has recently been completed, and utilized a number of behavioral tasks that probe multiple aspects of higher-order cognitive functions (35). Previous work by Thompson and colleagues on 4521 youth from the first release of baseline ABCD data distinguished three major higher-cognitive domains: General Ability, Speed/Flexibility, and Learning/Memory (36).

The present work links neurocognitive scores for these domains with resting state brain connectivity patterns. We first produced resting state connectomes for each subject, using multiple methods to control for the effects of head motion, a potentially serious confound in resting state studies (37). In a sample of 2,013 subjects who met quality control and other inclusion criteria, we next applied a recently developed multivariate predictive modeling method, brain basis set (BBS) (38, 39) (see Figure 1). This method takes advantage of the fact that though functional connectomes are large and complex, encompassing tens of thousands of connections, there is massive redundancy in the set of connections that differ across people. This allows a small set of components—we used 75 in the present study—to capture most meaningful inter-individual variation in connectomes (38, 39). We coupled BBS with leave-one-site-out cross-validation. That is, we train a BBS predictive model in all of the sites except one, apply the trained model to the held out site, and repeat this process with each site being held out. This allows us to gauge the generalizability of our predictive models to new, unseen subjects.

**Figure 1:**
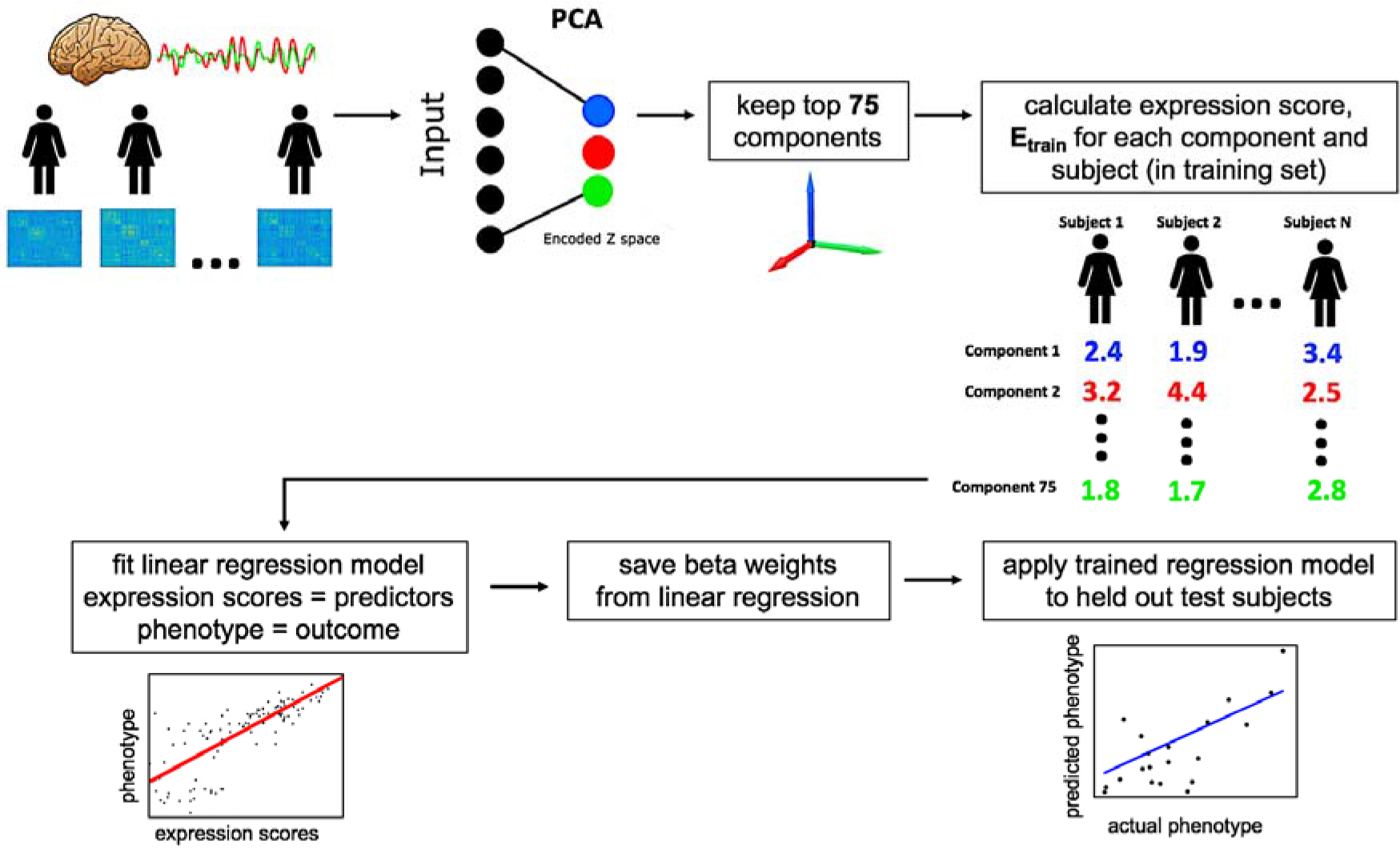
Main Steps of Brain Basis Set (BBS) Modeling. BBS is a multivariate predictive modeling method. It utilizes dimensionality reduction with principal components analysis (PCA) to construct a basis set for predicting phenotypes of interest.

### Resting state functional connectivity patterns are statistically significant predictors of neurocognitive scores for all three neurocognitive domains

We applied BBS to separately train predictive models for each of the three neurocognitive component scores. We then assessed the correlations between actual versus predicted neurocognitive scores, averaging across folds of the leave-one-site out cross-validation procedure. Results for each neurocognitive domain were: General Ability *r* = 0.31 (permutation *p* value < 0.0001, i.e., observed correlation was higher than all 10,000 correlations in the permutation distribution); Speed/Flexibility *r* = 0.06 (permutation *p* value = 0.02); Learning/Memory *r* = 0.15 (permutation *p* value < 0.0001, i.e., observed correlation was higher than all 10,000 correlations in the permutation distribution). Figure 2 displays consensus maps that highlight connections that were weighted more heavily in the respective predictive models.

**Figure 2:**
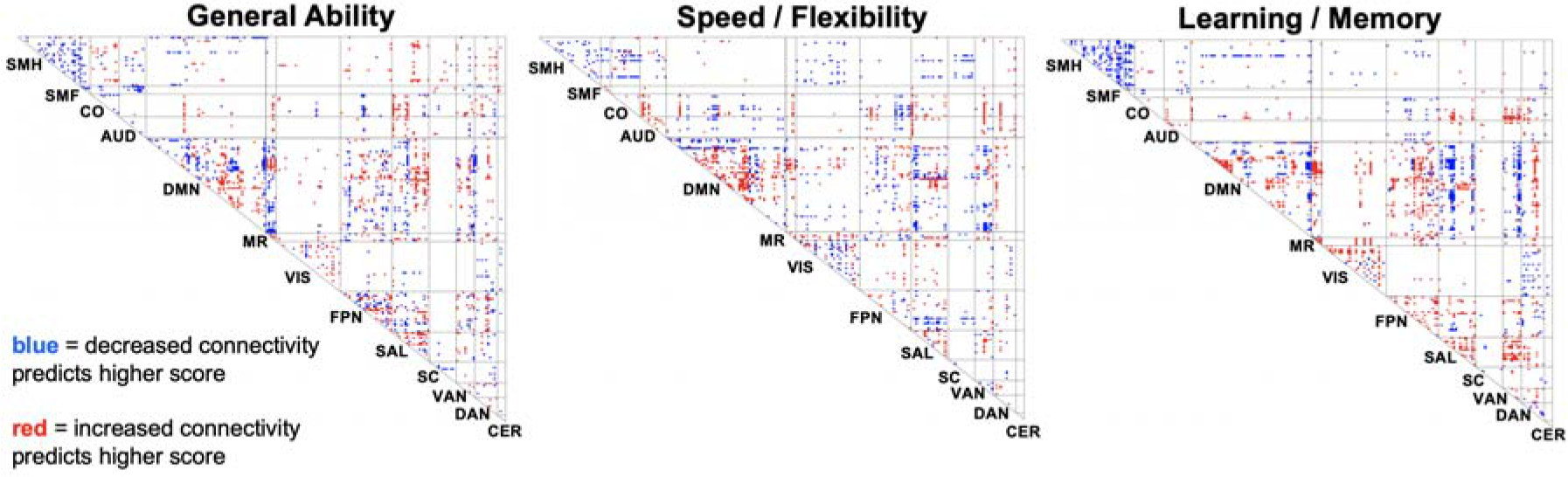
Consensus Maps of Connections that are Predictive of Neurocognitive Scores. Connections in task control networks and DMN are especially well represented among the set of connections predictive of neurocognitive scores. SMH=Somatomotor-hand, SMF=Somatomotor-faces, CO=Cingulo-opercular, AUD=Auditory, DMN= Default mode network, MR=Memory retrieval, VIS=Visual, FPN=Fronto-parietal, SAL=Salience, SC=Subcortical, VAN=Ventral Attention, DAN=Dorsal Attention, CER=Cerebellum.

In a secondary analysis, we included a number of covariates in these BBS models, including age, gender, race/ethnicity, highest parental education, household marital status, and household income. Correlations between actual versus predicted neurocognitive scores, averaging across folds of the cross-validation, were: General Ability *r* = 0.29 (permutation *p* value < 0.0001, i.e., observed correlation was higher than all 10,000 correlations in the permutation distribution); Speed/Flexibility *r* = 0.05 (permutation *p* value = 0.11); Learning/Memory *r* = 0.10 (permutation *p* value = 0.01).

### Predictive models for General Ability show evidence of substantial cross-site generalizability

Figure 3 shows the per-site results of the BBS-based predictive models. For General Ability, results were consistent across sites, with statistically significant correlations between predicted and actual scores achieved in 14 out of 15 held-out sites (all *p* values < 0.05). An important open question in neuroimaging is whether predictive models based on resting state connectivity patterns trained in one dataset can effectively generalize to unseen data, in particular when the unseen data is collected at a different site using a different MRI scanner. Results observed with General Ability scores demonstrate that successful generalization to data from unseen sites and scanners can indeed be achieved.

**Figure 3:**
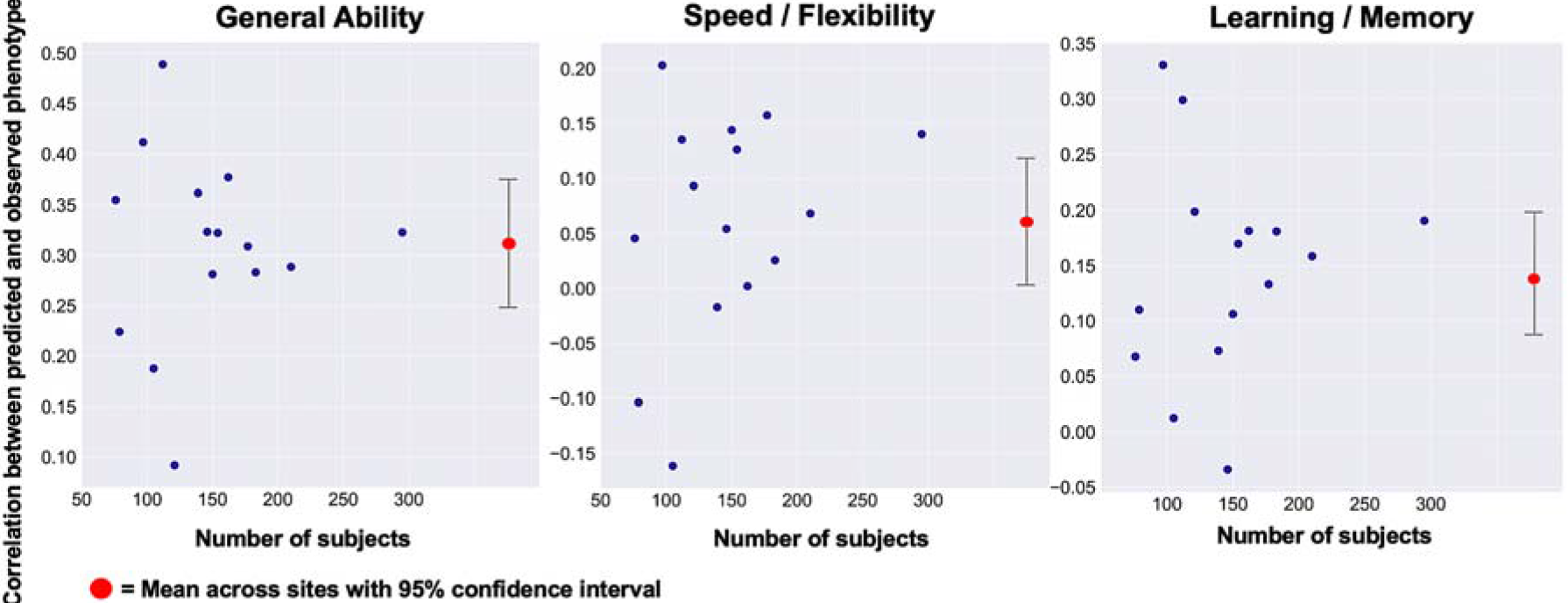
Per-Site Results of Predictive Models. We built multivariate models for predicting participants’ neurocognitive scores and tested these models with leave-one-site-out cross-validation across 15 sites. The General Ability neurocognitive score exhibited particularly strong generalization across sites.

### Connections within and between task control networks and DMN play a prominent role in prediction of neurocognitive scores

We use the term “task control-DMN intersection” to refer to: (i) connections within a task control network; (ii) connections within DMN, and (iii) connections that link a task control network node with a DMN node. Such connections represent roughly 20% of the total number of connections (6754 out of 34716 total) in the connectome. They represent, however, a much larger percentage of the suprathreshold connections in the consensus maps (Figure 2, where a *z*=2 threshold is used), in particular 43%, 43%, and 49% for General Ability, Speed/Flexibility, and Learning/Memory, respectively.

To further assess the importance of connections in the task control-DMN intersection, we dropped all connections outside this intersection and redid our BBS-based predictive modeling. The correlations between actual versus predicted neurocognitive scores, averaging across folds of the cross-validation, were: General Ability *r* = 0.26 (permutation *p* value < 0.0001, i.e., observed correlation was higher than all 10,000 correlations in the permutation distribution); Speed/Flexibility *r* = 0.07 (permutation *p* value = 0.02); Learning/Memory *r* = 0.08 (permutation *p* value = 0.002). In addition, we examined prediction of the three neurocognitive component scores when retaining pairs of networks (Figure 4). This analysis too suggested disproportionate importance of the task control-DMN intersection. In particular, DMN and task control networks emerged as three of the top four most important networks for all three neurocognitive factors.

**Figure 4:**
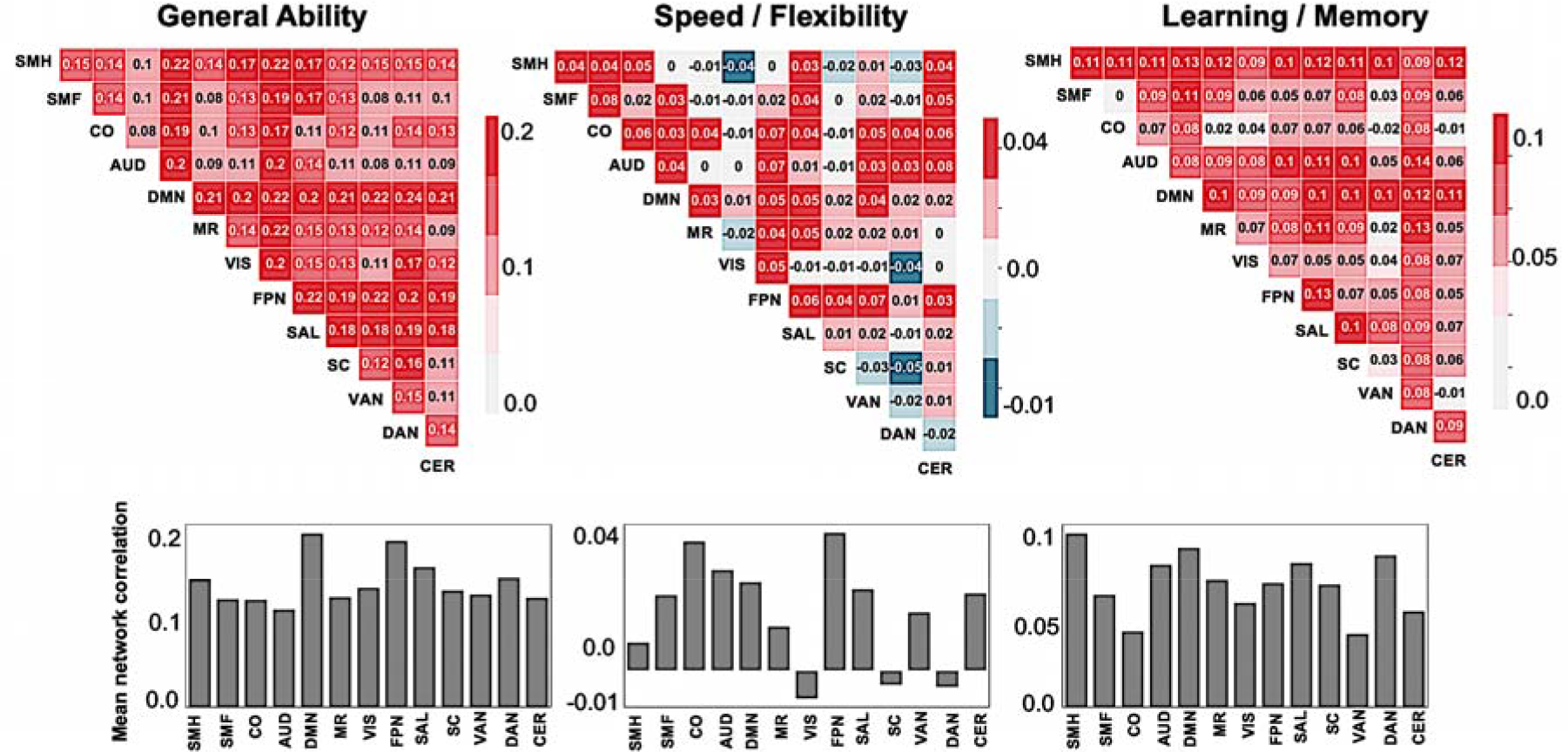
Prediction of Neurocognitive Domains with Two Networks: To assess the importance of networks for the success of the BBS predictive models, we trained new BBS models with just two networks (top row). We also calculated the mean performance for each network (bar graphs in bottom row). Values indicate correlations between actual and predicted scores, averaged across folds of the cross-validation analysis. These analyses revealed prominent roles for DMN and task control networks. SMH=Somatomotor-hand, SMF=Somatomotor-faces, CO=Cingulo-opercular, AUD=Auditory, DMN= Default mode network, MR=Memory retrieval, VIS=Visual, FPN=Fronto-parietal, SAL=Salience, SC=Subcortical, VAN=Ventral Attention, DAN=Dorsal Attention, CER=Cerebellum.

These findings highlighting the importance of the task control-DMN intersection for neurocognitive functioning can be interpreted in terms of recent theories of adaptive task control (16, 40). These accounts propose that task control networks modulate activity in distributed brain regions to facilitate cognitive control during complex tasks. In addition, a number of lines of evidence point to the DMN in particular as an important target of these top-down adaptive control signals. Modulation of the DMN is required in cases where task control networks must cooperate with DMN for coordinated processing, such as during complex problem-solving and prospective decision-making (23–25). It is also required to avoid interference by DMN (26) during externally focused, cognitively demanding tasks, with inadequate regulation of the DMN by task control networks sometimes leading to impaired task performance (41, 42).

### Connections predicting neurocognitive scores prominently include hub-like structures linking task control networks and DMN

The human brain exhibits a modular architecture in which most connections are within network (43, 44). Given this configuration, connector hubs that link two networks are critical for cross-network information flow (45, 46, 30). It has recently been argued that connector hubs in task control networks are important for propagating top-down adaptive control signals to distributed processing regions including the default network (16, 30).

Consistent with this idea, we noted hub-like structures that connect specific task control nodes to widespread regions of DMN in the consensus maps (in Figure 2, these hubs are seen as vertical “lines” in the DMN row). These hub-like structures appear especially prominent in the Learning/Memory consensus map, and we display the hubs for this consensus map in Figure 5.

**Figure 5:**
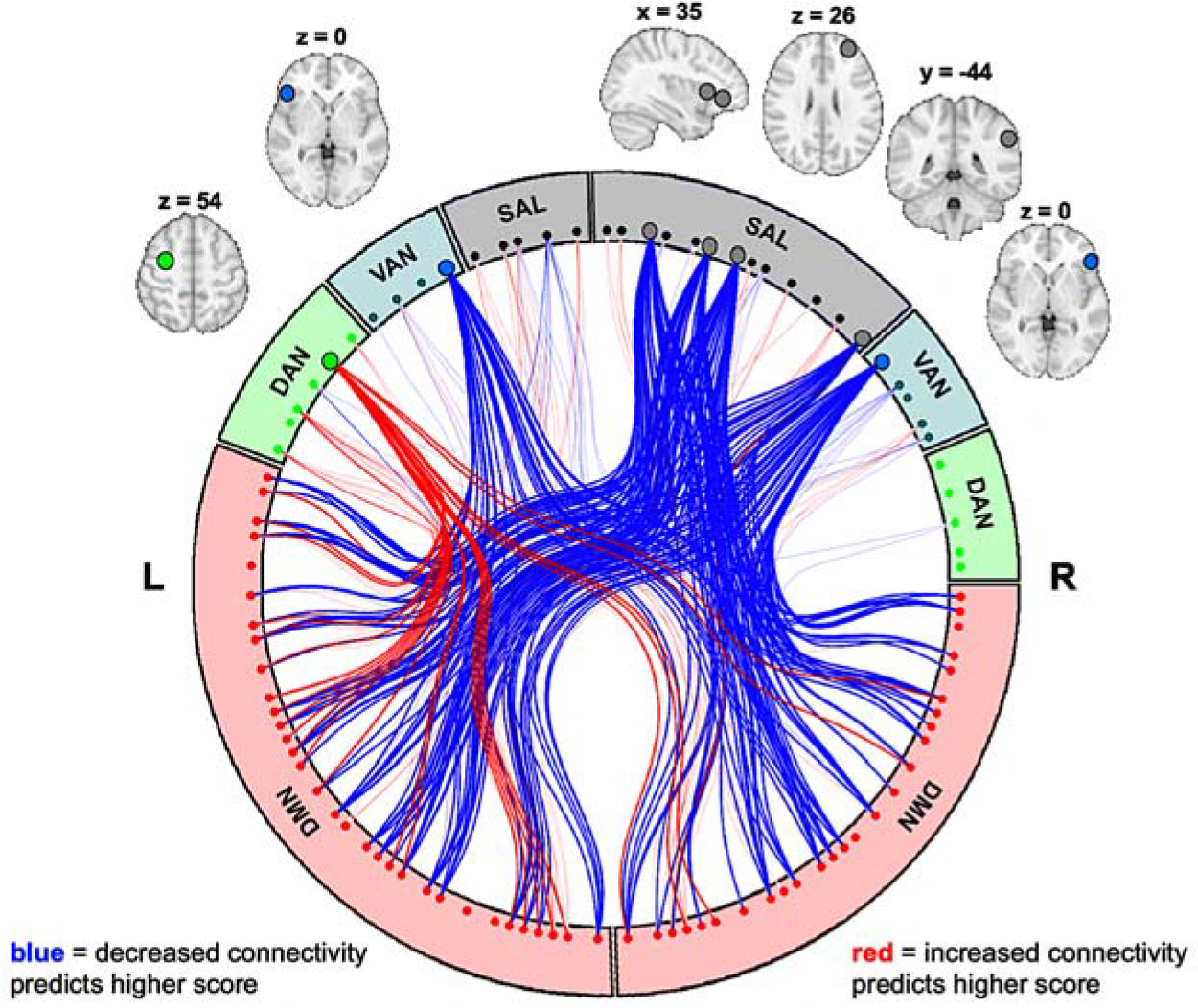
Hub-like Structures Linking Task-Control Networks and DMN. We observed hub-like structures in the consensus map for Learning/Memory in salience network (SAL), ventral attention network (VAN), and dorsal attention network (DAN). Each hub-like structure exhibited altered patterns of connectivity with nearly every node in default mode node network (DMN).

## Conclusions

In one of the largest neuroimaging investigations of youth to date, we demonstrate that resting state connectivity patterns are predictive of three major domains of higher-order cognitive functions: General Ability, Speed/Flexibility, and Learning/Memory. Our strongest results were observed with General Ability scores, where connectivity-based predictive models captured nearly 10% of the variance in scores and yielded statistically significant predictions in 14 out of 15 held-out sites. There is great interest in cognitive and psychiatric neuroscience in moving towards biomarkers, i.e., objective quantitative measures, of psychologically- or clinically- important constructs. Our results suggest that resting state brain connectivity patterns could provide one avenue for eventually producing effective, generalizable biomarkers of neurocognitive profiles in youth.

We also demonstrated that connections within and between task control networks and DMN are especially important for individual-differences in youth neurocognitive functioning across all three assessed domains. It is notable that connections involving task control networks and DMN are among the most vigorously maturing during youth (47, 33, 32, 48, 34), with pronounced increases in intra-DMN connectivity from early childhood to the mid-20’s (47, 34) and concurrent segregation of DMN from certain task control networks (33, 34). Given these observations, a major strength of the ABCD longitudinal design is that these same children will be followed and scanned at regular intervals through adolescence over the next 10 years. Thus, researchers accessing subsequent waves of ABCD data will be well positioned to produce detailed maturational trajectories of task control-DMN connections, a potentially critical locus of individual differences in neurocognitive profiles in health and disease.

## Methods

### 1. Sample and Data

The ABCD study is a multisite longitudinal study established to investigate how individual, family, and broader socio-cultural factors shape brain development and health outcomes. The study has recruited 11,875 children between 9-10 years of age from 21 sites across the United States for longitudinal assessment. At each assessment wave, children undergo assessments of neurocognition, physical health, and mental health, and also participate in structural and functional neuroimaging. Detailed description of recruitment procedures (49), assessments (50), and imaging protocols (51) are available elsewhere. The ABCD data repository grows and changes over time. The ABCD data used in this report came from NDA Study 576, DOI 10.15154/1412097, which can be found at https://ndar.nih.gov/study.html?id=576.

### 2. Data Acquisition, fMRI Preprocessing, and Connectome Generation

Imaging protocols were harmonized across sites and scanners. high spatial (2.4 mm isotropic) and temporal resolution (TR=800 ms) resting state fMRI was acquired in four separate runs (5min per run, 20 minutes total). For the current analysis, minimally preprocessed resting-state fMRI data from the curated ABCD annual release 1.1 were used, and full details are described in (52). This data reflects the application of the following steps: i) gradient-nonlinearity distortions and inhomogeneity correction for structural data; and ii) gradient-nonlinearity distortion correction, rigid realignment to adjust for motion, and field map correction for functional data. The data were subsequently further preprocessed by our group using SPM12. Anatomical (T1 weighted) images were co-registered to functional data. T1 weighted images were then registered linearly and nonlinearly to MNI space using the CAT12 Toolbox and DARTEL. This warp field was applied to the functional data to bring all subjects into template space and smoothed with a 6mm FWHM Gaussian kernel. Removal of artifacts arising from head motion was performed on the smoothed functional time series using ICA-AROMA (53).

The smoothed images then went through a number of resting state processing steps, including motion artifact removal steps comparable to the type B (i.e., recommended) stream of Siegel et al. (54). These steps include linear detrending, CompCor to extract and regress out the top 5 principal components of white matter and CSF (55), bandpass filtering from 0.1-0.01Hz, and motion scrubbing of frames that exceed a framewise displacement of 0.5mm.

We calculated spatially-averaged time series for each of 264 4.24mm radius ROIs from the parcellation of Power et al. (56). We then calculated Pearson’s correlation coefficients between each ROI. These were then were transformed using Fisher’s *r* to *z*-transformation.

### 3. Constructing Neurocognitive Component Scores

Component scores for three neurocognitive domains were derived through procedures described in detail in Thompson et al. (36), and presented here in brief. Thompson et al. conducted an exploratory analysis of ABCD neurocognitive assessments using Bayesian Probabilistic Principal Components Analysis (BPPCA; 80). This analysis used random effects for site and family to account for correlation among subjects in component scores and in residuals caused by the nested structure of the data. The model was implemented using the Bayesian inference engine stan (58) using the R package rstan to interface with R Version 3.4.291. This yielded a three component solution: (a) General Ability component (variance explained = 21.2%, [20.1%, 22.5%]) with strongest loadings for the NIH Toolbox Picture Vocabulary, Toolbox Oral Reading, and Little Man spatial reasoning tasks; (b) Speed/Flexibility component (20.5% [19.5%, 21.5%]) with strongest loadings from the NIH Toolbox Pattern Comparison Processing Speed task, NIH Toolbox Dimensional Change Card Sort, and the NIH Toolbox Flanker; and (c) Learning/Memory component (17.9% [16.9%, 19.1%]) with strongest loadings from the NIH Toolbox Picture Sequence Memory task and the Rey Auditory Verbal Learning Task (RAVLT) total number correct. In addition, the NIH Toolbox List Sort Working Memory task was represented on both the General Ability component and the Learning/Memory component. See Table S1 for details on factor loadings. We refer to these components as “neurocognitive components” throughout to distinguish them from the functional connectivity components that are used in the BBS predictive modeling approach (described below).

### 4. Inclusion/Exclusion

There were 4521 subjects in the ABCD Release 1.1 dataset. Of these, 3575 subjects had usable T1w images and one or more resting state runs that passed ABCD quality checking standards (fsqc_qc = 1). Next, 3544 passed preprocessing and were subsequently visually checked for registration and normalization quality, where 197 were excluded for poor quality. Motion was assessed based on number of frames censored, with a framewise displacement threshold of 0.5mm, and only subjects with two or more runs with at least 4 minutes of good data were included (*n*=2757). To remove unwanted sources of dependence in the dataset, only one sibling was randomly chosen to be retained for any family with more than 1 sibling (*n*=2494). Finally, in order to implement leave-one-site-out cross validation, sites with fewer than 75 subjects that passed these quality checks were dropped, leaving 2206 subjects across 15 sites to enter the PCA step of BBS predictive modeling. Thompson et al. (36) excluded 428 children due to incomplete neurocognitive data, and our prediction analyses was correspondingly restricted to only those subjects that had the three neurocognitive factors from their analysis. This left 2013 subjects across the 15 sites for the prediction step of BBS, and the demographic characteristics of this sample are shown in Table S2. For the analysis including more covariates (described in section 6) 1858 subjects were included due to additional missing data in the covariates.

### 5. Brain Basis Set Modeling (BBS)

BBS is a validated multivariate predictive method that uses dimensionality reduction to produce a basis set of components to make phenotypic predictions (see Figure 1 for an overview) (34, 38). For the dimensionality reduction step, we submitted an *n* subjects *x p* connections matrix from a training dataset for principal components analysis using the pca function in MATLAB, yielding *n*-1 components ordered by descending eigenvalues (note that that p > n). We select the top 75 components for our basis set based on our previous work showing that somewhere between 50 to 100 components yields optimal prediction of a broad array of behavioral phenotypes (38, 39), with inclusion of additional components typically reducing performance due to overfitting.

Next, in the training dataset, we calculate the expression scores for each of the 75 components for each subject by projecting each subject’s connectivity matrix onto each principal component. We then fit a linear regression model with these expression scores as predictors and the phenotype of interest as the outcome, saving **B**, the 75 *x 1* vector of fitted coefficients, for later use. In a test dataset, we again calculate the expression scores for each of the 75 components for each subject. Our predicted phenotype for each test subject is the dot product of **B** learned from the training dataset with the vector of component expression scores for that subject.

### 6. Leave-One-Site-Out Cross Validation

To assess of the performance of BBS-based prediction models, we used leave-one-site-out cross-validation, which was performed separately for each of the three neurocognitive component scores. In each fold of the cross-validation, data from one of the 15 sites served as the held-out test dataset and data from the other 14 sites served as the training dataset. Additionally, at each fold of the cross-validation, we did the following: 1) PCA was performed on the training dataset yielding a 75-component basis set; 2) a BBS model was trained to predict the relevant neurocognitive factor as the outcome variable. These BBS models included covariates for head motion (mean FD and mean FD squared), and in applying these trained BBS models to the held-out test dataset, the level of these covariates was set at zero. In a secondary analysis, we used a more extensive set of covariates in these BBS predictive models, including age, gender, race/ethnicity, highest parental education, household marital status, and household income.

### 7. Permutation Testing

Cross-validation, as opposed to validation in a completely independent dataset, is associated with elevated variance of estimates (59). Thus we assessed the significance of all cross-validation-based correlations with non-parametric permutation tests (60).

The distribution under chance of correlations between BBS-based predictions of neurocognitive scores and observed neurocognitive scores was generated by randomly permuting the 2013 subjects’ neurocognitive scores 10,000 times. At each iteration, we performed the leave-one-site out cross validation procedure described above (which includes refitting BBS models at each fold of the cross-validation). We then recalculated the average correlation across folds between predicted versus actual neurocognitive scores. The average correlation across folds that was actually observed was located in this null distribution in terms of rank, and statistical significance was set as this rank value divided by 10,000.

Since the BBS models fit at each iteration of the permutation test included covariates (mean FD and mean FD squared for the main model; additional covariates in the secondary model), the procedure of Freedman and Lane was followed (61). In brief, a BBS model was first estimated with nuisance covariates alone, residuals were formed and were permuted. The covariate effects were then added to the permuted residuals, creating an approximate realization of data under the null hypothesis, and the statistical test of interest was calculated on this data (see FSL Randomise http://fsl.fmrib.ox.ac.uk/fsl/fslwiki/Randomise/Theory for a neuroimaging implementation).

### 8. Consensus Component Maps for Visualization

We used BBS with 75 whole-connectome components to make predictions about neurocognitive component scores. To help convey overall patterns across the entire BBS predictive model, we constructed “consensus” component maps. We first fit a BBS model to the entire dataset consisting of all participants across the 15 included sites. We then multiplied each component map with its associated beta from this fitted BBS model. Next, we summed across all 75 components yielding a single map, and *z* scored the entries at *z*=2. The resulting map indicates the extent to which each connection is positively (red) or negatively (blue) related to the covariate of interest.

## Supporting information

## Acknowledgments

Data used in the preparation of this article were obtained from the Adolescent Brain Cognitive Development (ABCD) Study (https://abcdstudy.org), held in the NIMH Data Archive (NDA). This is a multisite, longitudinal study designed to recruit more than 10,000 children age 9-10 and follow them over 10 years into early adulthood. The ABCD Study is supported by the National Institutes of Health and additional federal partners under award numbers U01DA041022, U01DA041028, U01DA041048, U01DA041089, U01DA041106, U01DA041117, U01DA041120, U01DA041134, U01DA041148, U01DA041156, U01DA041174, U24DA041123, U24DA041147, U01DA041093, and U01DA041025. A full list of supporters is available at https://abcdstudy.org/federal-partners.html. A listing of participating sites and a complete listing of the study investigators can be found at https://abcdstudy.org/Consortium_Members.pdf. ABCD consortium investigators designed and implemented the study and/or provided data but did not necessarily participate in analysis or writing of this report. This manuscript reflects the views of the authors and may not reflect the opinions or views of the NIH or ABCD consortium investigators.

This work was supported by the following grants from the United States National Institutes of Health, the National Institute on Drug Abuse, and the National Institute on Alcohol Abuse and Alcoholism: R01MH107741 (CS), U01DA041106 (CS, LH, MH), 1U24DA041123-01 (WT), U01DA041120 (ML), T32 AA007477 (AW). In addition, CS was supported by a grant from the Dana Foundation David Mahoney Neuroimaging Program. This research was supported in part through computational resources and services provided by Advanced Research Computing at the University of Michigan, Ann Arbor.

## Author Contributions

Conceptualization: CS, MA, SR; Methodology: CS, MA, SR; Formal Analysis: CS, MA, ML, SR, WT; Data Curation: MA, SR; Writing – Original Draft: CS; Writing – Reviewing and Editing; AW, CS, LH, MA, MH, ML, SR, WT; Visualization: MA, SR; Supervision: CS, MH; Funding Acquisition: CS, MH.

