## Supplementary material for "Prediction of Neurocognitive Profiles in Youth From Resting State fMRI"

**Supplement**

**Supplementary Tables**

| Test | General Ability | Speed/Flexibility | Learning/Memory |
| --- | --- | --- | --- |
| Pic Vocab | 0.754 [0.706, 0.799] | 0.065 [0.029, 0.102] | 0.190 [0.133, 0.252] |
| Flanker | 0.213 [0.161, 0.26] | 0.712 [0.668, 0.754] | 0.067 [0.013, 0.119] |
| List | 0.471 [0.4, 0.538] | 0.148 [0.105, 0.195] | 0.493 [0.416, 0.563] |
| Card Sort | 0.205 [0.163, 0.252] | 0.710 [0.668, 0.751] | 0.232 [0.184, 0.287] |
| Pattern | 0.015 [-0.029, 0.055] | 0.813 [0.771, 0.85] | 0.085 [0.039, 0.135] |
| Picture | 0.012 [-0.023, 0.049] | 0.135 [0.102, 0.171] | 0.863 [0.816, 0.904] |
| Reading | 0.820 [0.782, 0.86] | 0.123 [0.084, 0.16] | 0.122 [0.067, 0.173] |
| RAVLT | 0.306 [0.253, 0.364] | 0.125 [0.085, 0.163] | 0.712 [0.663, 0.76] |
| LMT | 0.500 [0.424, 0.57] | 0.299 [0.246, 0.36] | 0.068 [0.002, 0.144] |

**Table S1**. **Neurocognitive Factor Loadings**. Values of factor loadings from the Bayesian Probabilistic PCA (following Thompson et al. (1)). Numbers outside of the brackets represent the posterior median, which is the most likely value of each loading. Numbers reported in brackets represent 95% posterior credible intervals for the values, which indicate that there is a 95% chance that the true loading falls within this range. Pic Vocab = Toolbox Picture Vocabulary; Flanker = Toolbox List Sort Working Memory Task; Card Sort = Dimensional Change Card Sort Task; Pattern = Toolbox Pattern Comparison Processing Speed Task; Picture = Toolbox Picture Sequence Memory Task; Reading = Toolbox Oral Reading Test; RAVLT = Rey Auditory Verbal Learning Task, total correct on learning trials; LMT = Little Man Task proportion correct.

|  | Included | Excluded | P-value |
| --- | --- | --- | --- |
| N | 2206 | 2315 |  |
| Age (mean (s.d.)) | 10.02 (0.61) | 9.99 (0.61) | 0.08 |
| Female (%) | 1058 (48.0) | 1088 (47.0) | 0.57 |
| Race Ethnicity (%) |  |  | 8.03x10^-5^ |
| White | 1288 (58.4) | 1363 (58.9) |  |
| Black | 173 (7.84) | 271 (11.7) |  |
| Hispanic | 472 (21.4) | 415 (17.9) |  |
| Asian | 49 (2.22) | 54 (2.33) |  |
| Other | 218 (9.88) | 209 (9.03) |  |
| Unknown | 6 (0.272) | 3 (0.130) |  |
| Highest Parental Education (%) |  |  | 0.03 |
| < HS Diploma | 80 (3.63) | 98 (4.23) |  |
| Bachelor | 587 (26.6) | 635 (27.4) |  |
| HS Diploma/GED | 141(6.39) | 184 (7.95) |  |
| Post Graduate Degree | 807 (36.6) | 861 (37.2) |  |
| Some College | 589 (26.7) | 534 (23.1) |  |
| Household Marital Status (%) | 1571 (71.2) | 1633 (70.5) | 0.58 |
| Household Income (%) |  |  | 0.24 |
| <50K | 499 (22.6) | 452 (19.5) |  |
| >=100k | 914 (41.4) | 947 (40.9) |  |
| >=50k & <100K | 638 (28.9) | 625 (27.0) |  |
| Unknown | 0 (0.0) | 356 (15.4) |  |

**Table S2**. **Participant Demographics**. Demographic characteristics of full sample and sample included in the present analysis.
